# Replication fork plasticity is a therapeutic vulnerability in acute myeloid leukemia

**DOI:** 10.64898/2026.08.13.744634

**Authors:** Cyril Dördelmann, Tsz Kan Fung, Tristan Gasparetto, Larissa Mendes Bomfim, Chi Wai Eric So, Massimo Lopes

## Abstract

Uncontrolled proliferation of myeloid progenitor cells in acute myeloid leukemia (AML) is counteracted in most patients by toxic and often ineffective systemic treatments. Poly (ADP-ribose) polymerase inhibitors (PARPi) show subtype-restricted activity – potent in RUNX1-RUNX1T1 and PML-RARɑ fusions, limited in KMT2A-rearranged (KMT2A-r) disease – but the lack of molecular understanding has hampered their clinical implementation. We combined single-cell and single-molecule assays on DNA replication intermediates and DNA damage signalling with therapy response readouts to investigate the role of fork plasticity factors in response to PARPi and AML standard-of-care (cytarabine, araC). In PARPi-sensitive AML models, PARP inhibition deregulates RECQ1-mediated fork restart, initially triggering fork acceleration and later fork breakage within the same S phase. Conversely, PARPi resistant KMT2A-r AML lines are protected by PrimPol-dependent DNA synthesis and its inactivation promptly induces fork breakage and PARPi sensitivity. Strikingly, PrimPol overexpression in PARPi-sensitive AML models prevents fork collapse and PARPi/araC therapy response, both *in vitro* and *in vivo*, identifying PrimPol as novel predictive biomarker and therapeutic target in AML. Our data uncover novel tissue-specific mechanisms of action for PARPi and pinpoint replication fork plasticity as key molecular determinant of AML therapy response.

**Highlights:**

- Fork plasticity is a key molecular determinant of treatment response in leukemia.
- PARP inhibition triggers fork breakage via deregulated restart of reversed forks.
- Bypassing fork reversal, PrimPol limits therapy-induced DNA damage and cytotoxicity in AML.
- PrimPol drives resistance to cytarabine and PARP inhibition *in vitro* and *in vivo*.

## Introduction

Acute myeloid leukemia (AML) is a highly heterogeneous disease with a relatively low mutational burden compared with many solid tumors (Martincorena & Campbell, 2015). Recurrent, prognostically relevant mutations and oncofusions that affect transcriptional and epigenetic regulators are well defined in the field (Khoury et al., 2022; Zeisig et al., 2011). Although a few agents have recently been approved for molecularly defined AML subsets, classical chemotherapy – i.e. “7+3” induction via cytarabine (araC) plus an anthracycline – remains standard-of-care in most cases and frequently proves toxic or ineffective, especially in elderly patients (Kantarjian et al., 2025). AML targeting strategies that exploit replication stress and DNA repair – most prominently poly (ADP-ribose) polymerases inhibitors (PARP inhibitors; PARPi) and polymerase-theta (POLQ/Polθ) inhibitors – have recently attracted interest for clinical use. The main obstacles to their final approval are the toxic side-effects in a subset of patients and the lack of molecular biomarkers for patient stratification in clinical trials (Padella et al., 2022; Vekariya et al., 2023). Given the unmet clinical need to improve AML therapeutic options, uncovering the molecular determinants of AML response to PARPi and standard-of-care is strictly needed to reduce undesired side effects and to select patients that will likely benefit of established and novel therapeutic regimens.

Replication stress (RS) is a hallmark driver of genome instability (Macheret & Halazonetis, 2015) that is well documented in established AML and increasingly recognized as an early event in leukemogenesis (Flach et al., 2014; Kwok et al., 2024). Endogenous genotoxins – reactive oxygen species (ROS) and aldehydes – together with AML lineage-associated mutations (e.g., DDX41, SF3B1) impose replication stress in hematopoietic stem and progenitor cells (HSPCs), promoting genomic instability and malignant transformation (Dingler et al., 2020; Flach et al., 2014; Lee et al., 2025; Pontel et al., 2015; Rombaut et al., 2024; Sallmyr et al., 2008).

Nuclear PARPs (PARP1-3) safeguard genome integrity by coordinating base-excision and single-strand break repair (BER/SSB), Okazaki-fragment maturation, replication-fork stability/restart, double-strand break repair and chromatin remodelling (Azarm & Smith, 2020). PARPi generally act through catalytic inhibition as well as drug stabilized PARP-DNA trapping, converting non-lethal fork perturbations into ssDNA gaps, transcription-replication conflicts and fork collapse. These lesion and obstacles are particularly toxic in a subset of solid tumours that are defective in homologous recombination (HR), thereby affecting fork stability and break repair. Unfortunately, resistance commonly arises and was proposed to reflect HR restoration, fork-stabilization rewiring, trapping alterations, drug efflux and/or post-replicative gap suppression by Polθ or REV1/Polζ translesion synthesis (TLS) (Dibitetto et al., 2024; Kim & Yu, 2025; Rose et al., 2020; Zou et al., 2025). Durable responses were so far mostly reported in HR-deficient ovarian cancer patients, while modest gains in progression-free survival are observed in germline BRCA1/2-mutated pancreas cancer patients (Hage Chehade et al., 2025). However, the detailed molecular mechanisms mediating PARPi toxicity in solid tumours remain debated and are even more elusive in leukemia, where a recognizable HR-defective subset is lacking.

Replication forks facing endogenous or exogenous obstacles frequently undergo fork reversal via complex biochemical reactions involving the central recombinase RAD51 and specialised DNA translocases (i.e. HLTF, SMARCAL1 and ZRANB3), requiring fork restart by the specialised helicase RECQ1. Such complex and slow remodelling/restart of replication forks is counterbalanced by an alternative and faster damage-bypass strategy; this involves PrimPol-mediated repriming and leaves post-replicative ssDNA gaps that will eventually need repair. The balance between remodelling and repriming is finely tuned in solid cancer cells and affects the cellular response to chemotherapeutic treatments interfering with the replication process (Bai et al., 2020; Quinet et al., 2020). Intriguingly, PARP inhibition was proposed to boost reversed-fork restart and to induce accelerated fork progression, which by itself may cause replication stress and genome instability (Berti et al., 2013; Calzetta et al., 2025; Maya-Mendoza et al., 2018; Zellweger et al., 2015). However, current models of PARPi toxicity still mostly focus on SSB/DSB induction and repair, while the contribution of fork acceleration, remodelling and restart in PARPi toxicity has not been thoroughly explored.

Recent studies have uncovered how different tissues physiologically adopt distinct strategies of replication fork plasticity, especially when responding to proliferation stimuli. Epidermis and germinal-centre B cells often rely on TLS for lesion bypass, whereas fast-cycling compartments – e.g. embryonic stem cells and HSPCs – exhibit opposite adaptations in replication fork speed and replication stress response. Despite their rapid cell cycles, embryonic stem cells replicate with comparatively slow forks (Ahuja et al., 2016; Delbos et al., 2005; Jacobs et al., 2022; Kushinsky et al., 2024; Schuch et al., 2017; Ubieto-Capella et al., 2024). Conversely, HSPCs drastically accelerate replication fork progression during stress-induced proliferation via PrimPol-dependent repriming, supporting expansion and engraftment (Jacobs et al., 2022). In AML, DNA-damage response (DDR) programs are also deregulated and differ by subtype. PARPi resistant KMT2A-rearranged AML often retain DDR proficiency, while PARPi sensitive PML-RARɑ and RUNX1-RUNX1T1 downregulate DDR pathways (Esposito et al., 2015; Park et al., 2021; Zhao et al., 2016; Zhao & So, 2017). However, a causative link between these transcriptional effects and PARPi response in AML has not been established. In fact, these unresolved mechanistic questions have limited the rationale design of PARPi clinical trials in these tumours.

Here, we hypothesized that specific differences in replication fork progression and plasticity among AML subtypes may contribute to the differential drug response observed in AML. Combining single-cell and single-molecule assays, we show that AML cells of all tested subtypes exhibit accelerated replication forks. However, PARP inhibition specifically induces S-phase specific DNA damage in RUNX1-RUNX1T1 and PML-RARɑ AML subtypes, via deregulated RECQ1-dependent fork restart. In contrast, KMT2A-r AML upregulate *PRIMPOL* and engage this protein at forks to efficiently bypass fork reversal, thereby conferring resistance to PARP inhibitors and standard-of-care (i.e. araC). Moreover, high *PRIMPOL* expression correlates with poorer patient survival, markedly limits PARPi/araC cytotoxicity *in vitro* and impairs PARPi/araC treatment success in AML mouse models, identifying PrimPol as a promising biomarker and potential novel therapeutic target in otherwise chemoresistant AML.

## Results

### Fast fork progression in AML is vulnerable to further PARPi-induced acceleration

Upon physiological, stimuli-induced proliferation, hematopoietic stem and progenitor cells (HSPCs) markedly increase their replication speed (Jacobs et al., 2022). We hypothesized that accelerated fork speed may represent an adaptation and therefore a vulnerability of leukemic hyper-proliferating HSPCs. To test this hypothesis, we compared short-term cultured CD34+-enriched healthy human donor peripheral blood (HD PB) and bone marrow (HD BM) samples to four common AML cell lines Kasumi-1 (RUNX1-RUNX1T1), NB4 (PML-RARα), THP-1 and MOLM-13 (KMT2A-MLLT3). To measure replication fork speed at single-molecule level, cells were pulse-labelled with nucleotide analogues and replicated tracks were measured on spread DNA fibers (Jackson & Pombo, 1998). Similarly to stress-induced HSPCs (Jacobs et al., 2022), all AML cell lines display accelerated replication fork progression, compared to HD PB and BM samples (Figure 1A). Similar trends were observed comparing healthy Ckit+ mouse cells with isogenic cells expressing the corresponding leukemia-associated transcription factors (LATFs) Ae-9a, PML-RARα, KMT2A-MLLT3 upon retroviral transduction/transformation (Figure S1A; Zeisig & So., 2009). Along with previous observations in pre-leukemic disease (Wildschut et al., 2023), this evidence suggests that acceleration of fork progression could be a key molecular event during leukemic transformation.

**Figure 1.**
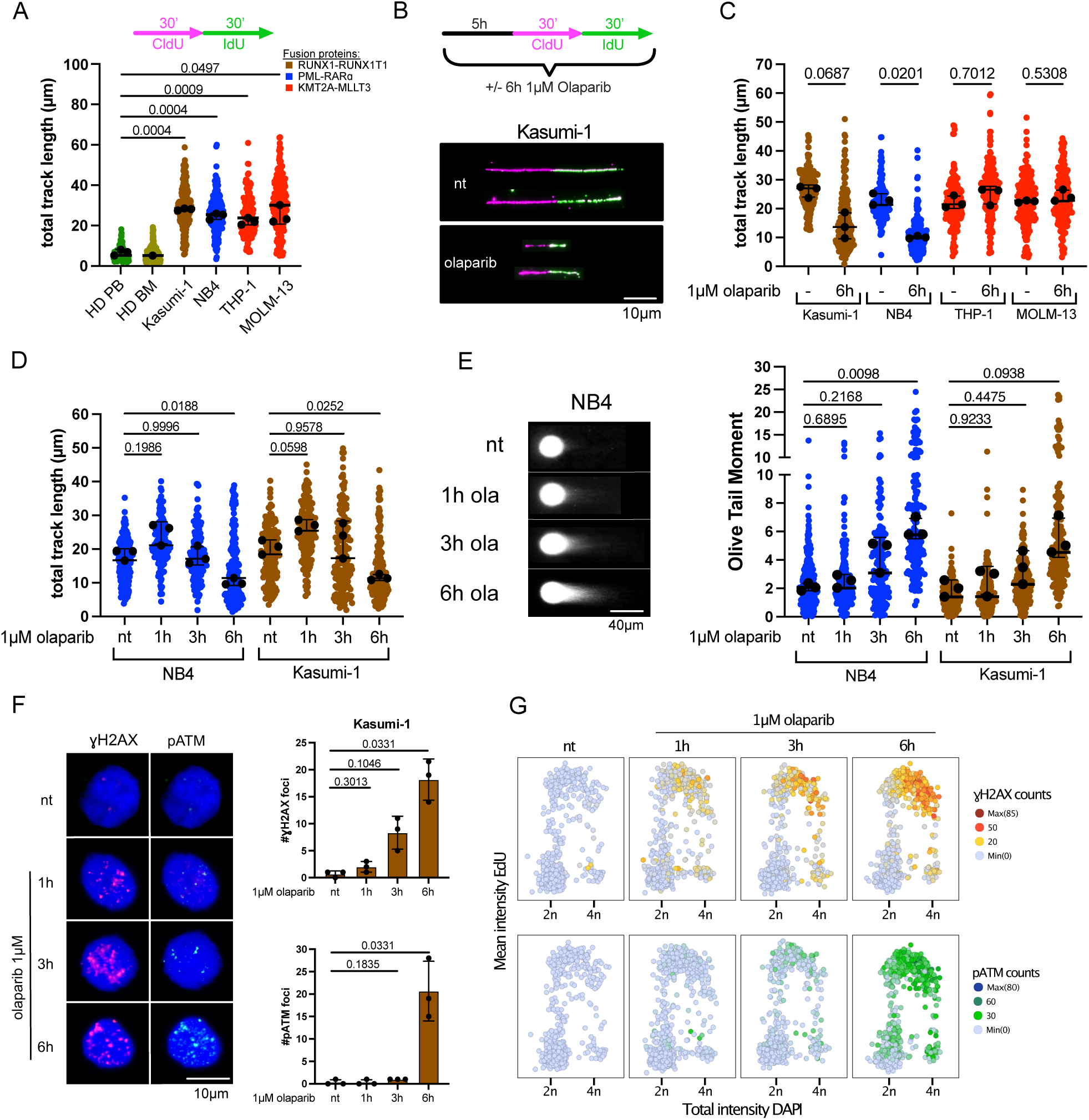
Fast replication fork progression in AML is vulnerable to further acceleration induced by PARP inhibition. (A) Labelling scheme and distribution of total CldU+IdU track lengths (ongoing forks) in CD34+ cells from healthy donors - peripheral blood (PB) or bone marrow (BM) – and human AML cell lines Kasumi-1 (*RUNX1-RUNX1T1*), NB4 (*PML-RARɑ*), THP-1 (*KMT2A-MLLT3*), MOLM-13 (*KMT2A-MLLT3*). ≥100 fibers were scored per condition per replicate. The horizontal black line indicate the median per condition. Overlaid black dots represent the per-experiment median from n = 3 independent experiments; error bars = SD of those medians. (B) DNA fiber assay schematic (top) and representative DNA fiber tracks (bottom) of Kasumi-1 cells. Cells were optionally treated with 1µM olaparib for 6 h; during the final hour, cells were sequentially labelled with CldU and IdU nucleotides as indicated. Immunostaining shows CldU (magenta) and IdU (green). To assess replication fork speed, double-labelled CldU/IdU tracts (ongoing forks) were measured. Scalebar, 10 µm0128 (C) Total fiber track lengths of a representative biological replicate of AML cell lines, untreated (-) or treated with 1µM olaparib for 6h. Black line marks the median. Overlaid black dots represent the per-experiment median from n = 3 independent experiments; error bars = SD of those medians. (D) Representative timecourse fiber replicate of total fiber track lengths in Kasumi-1 and NB4 cells untreated or 1, 3 and 6 h after 1µM of olaparib treatment. ≥100 fibers were scored per condition per replicate. Black line marks the median. Overlaid black dots represent the per-experiment medians (n = 3) ± SD (E) Neutral comet assay representative images (left) and Olive Tail Moment (OTM) measurements (right) in Kasumi-1 and NB4 of one representative biological replicate. Cells were untreated or 1, 3 and 6 h after 1µM of olaparib (Ola) treatment. ≥100 nuclei were scored per condition per replicate. Black line marks the median. Overlaid black dots represent the per-experiment medians (n = 3) ± SD. Scalebar, 40 µm (F) Representative images (left) and S-phase nuclear foci counts of ɣH2AX (Ser139) and pATM (S1981) immunofluorescence staining (right) in Kasumi-1 cells. Cells were untreated or 1, 3 and 6 h after 1µM olaparib treatment. 4’,6-diamidino-2-phenylindole (DAPI) is displayed to identify nuclei and ɣH2AX/pATM-negative cells. ≥200 cells were quantified per condition per replicate. Black dots represent the per-experiment medians (n = 3). Bars represent mean ± SD. Scalebar, 10 µm. (G) Quantitative image-based cytometry (QIBC) cell-cycle profiles of Kasumi-1 cells, plotted by DNA content (total DAPI intensity) and DNA synthesis (log-mean EdU intensity). Cells were untreated or 1, 3 and 6 h after 1µM olaparib treatment. Per-cell ɣH2AX (top) and pATM (bottom) foci counts are encoded by color. (F) Unpaired Welch’s t-test (A,B,D-F) Brown-Forsythe and Welch’s one-way ANOVA with Dunnett’s T3 multiple comparisons test. Indicated comparisons display numerical p-values.

Previous studies reported epigenetic deregulation of key replication stress response and DNA repair factors in AML (Esposito et al., 2015; Wong & So, 2020; Zeisig & So, 2021) and suggested that these may underlie the differential sensitivity of AML subtypes to poly (ADP-ribose) polymerase (PARP) inhibitors (PARPi) (Esposito et al., 2015). Besides various mechanisms recently proposed for PARPi cytotoxicity in different cell types (Dibitetto et al., 2024; Rose et al., 2020), PARP inhibition was recently linked to reversed replication fork restart and accelerated fork progression (Berti et al., 2013; Maya-Mendoza et al., 2018; Zellweger et al., 2015). Given the high rate of fork progression in untreated AML cells, we set out to investigate whether deregulated fork progression may contribute to PARPi toxicity in different AML subtypes. Short (6h) treatment with the PARP1/2 inhibitor olaparib at a dose of 1µM – which shows differential cytotoxicity in AML subtypes (Esposito et al., 2015) – triggered a drastic shortening of replicative tracts in two PARPi-sensitive cell lines (NB4, Kasumi-1), but not in the two KMT2A-rearranged cell lines (THP-1, MOLM-13), which are reportedly PARPi-resistant (Figure 1B,C; Esposito et al., 2015). These drastic effects of PARPi on replicated track length could be reproduced in mouse leukemia cell lines transformed with the corresponding fusion proteins (Figure S1B). Hence olaparib sensitivity is associated with PARPi effect on fork progression, suggesting that PARP inhibition may affect AML cells via replication-associated DNA damage.

To further investigate PARPi effects on fork acceleration and DNA damage accumulation, we conducted on the same cell populations kinetic DNA fiber assays and neutral DNA comet assays, to measure double-strand break (DSB) formation, as exemplified by ɣ-irradiation (Figure S1C). One hour after olaparib addition, both Kasumi-1 and NB4 cells displayed marked acceleration of replication forks without DNA damage accumulation. By three to six hours post-treatment, however, replication tracks progressively shortened, and the cells displayed increasing levels of DSBs (Figure 1D,E), strongly suggesting that replication fork acceleration by PARP1 inhibition precedes and triggers fork collapse in the PARPi-sensitive AML subtypes.

To further assess replication-associated DSBs, we employed immunofluorescence staining of DNA damage response (DDR) markers. Upon DSBs, H2AX histone tail phosphorylation at Ser139 (ɣH2AX) promptly occurs, and colocalizes with autophosphorylated (Ser1981) Ataxia Telangiectasia mutated (pATM) (Dar et al., 2006; Ewald et al., 2007; Sordet et al., 2009). Combining these DDR markers with DAPI counterstaining and pulse labelling of nascent DNA using 5-ethynyl-2’-deoxyuridine (EdU), we monitored DSB occurrence with cell-cycle resolution, using quantitative image-based cytometry (QIBC) (Toledo et al., 2013) (Figure S1D). Using this experimental setup to investigate PARPi response in AML, we found that prolonged olaparib exposure in Kasumi-1 led to rapid and gradual increase in S-phase-specific ɣH2AX foci, while elevated pATM foci were detected only after six hours (Figure 1F,G). This temporal separation suggests that the early increase in ɣH2AX predominantly reflect replication stress signalling lesions that are subsequently converted into replication-associated DNA breaks within the same S phase, eliciting ATM activation. At this later timepoint, ɣH2AX and pATM foci frequently co-localized (Figure S1E). By contrast, PARPi-resistant THP-1 cells showed only minimal increases in either marker (Figure S1F).

Altogether, these data indicate that, in PARPi-sensitive AML subtypes, olaparib further accelerates fast replication fork progression and thereby induces S-phase-specific DNA damage and DSB signalling.

### PARPi-mediated fork collapse in AML reflects deregulated fork restart by RECQ1

Fork acceleration by PARP inhibition was previously linked to multiple deregulations, including untimely activation of the reversed-fork restart helicase RECQ1 (Berti et al., 2013; Maya-Mendoza et al., 2018). To test whether RECQ1-mediated fork restart underlies the replication fork acceleration and genome instability in PARPi-sensitive AML subtypes, we established stable *shRECQ1* Kasumi-1 and NB4 cell lines (Figure 2A). Remarkably, RECQ1 depletion fully rescued both the severe tract shortening and the DNA breakage induced by 6h olaparib treatment (Figure 2B,C). In addition, RECQ1 depletion significantly improved viability after five days of continuous olaparib exposure in NB4 cells (Figure 2D).

**Figure 2.**
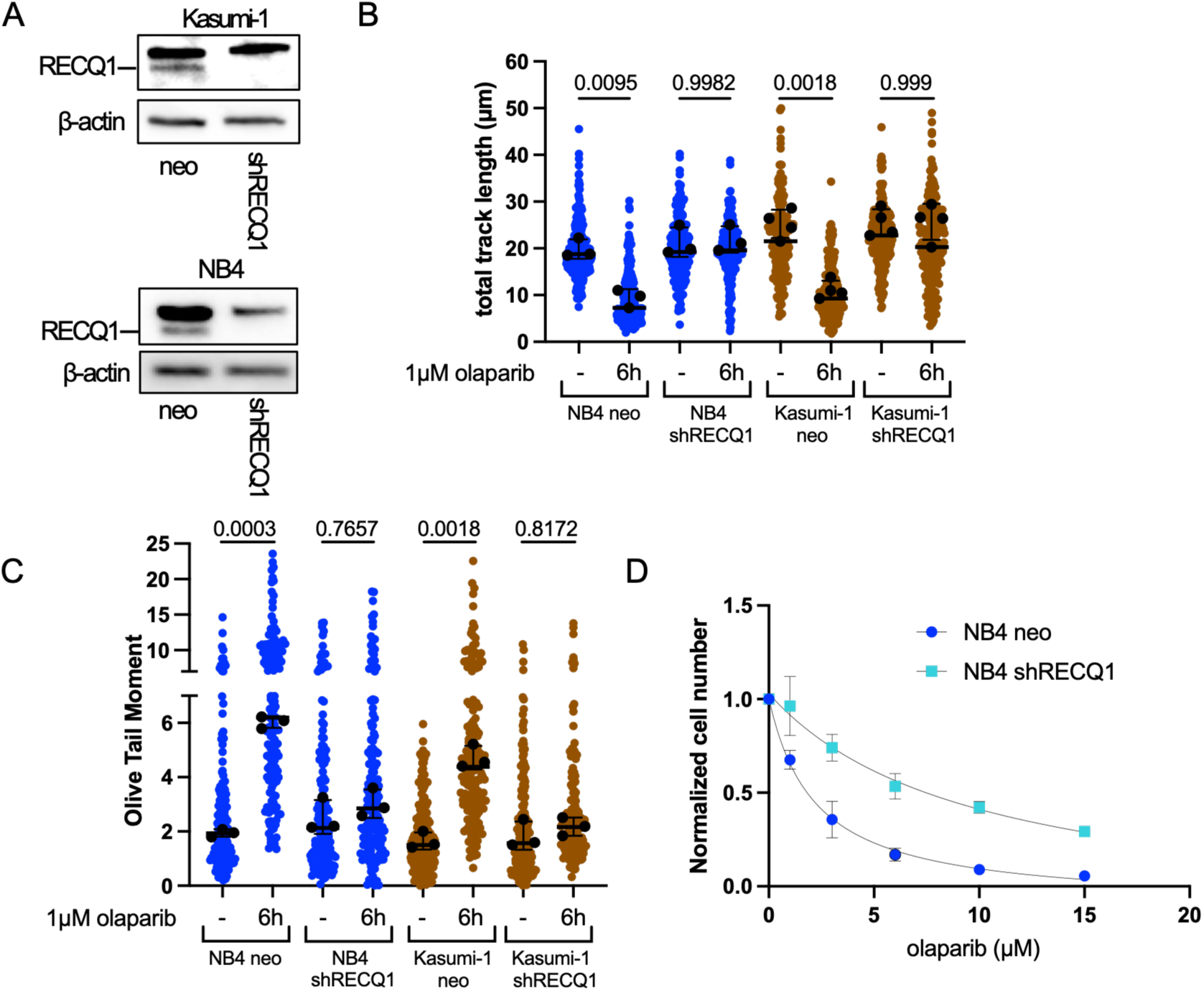
PARPi-mediated fork collapse in AML reflects deregulated fork restart by RECQ1. (A) RECQ1 (RECQL) immunblot in Kasumi-1 (top) and NB4 (bottom) versus their stable shRECQ1 counterparts showing efficient depletion. β-actin serves as loading control (B) Total fiber track length for a representative biological replicate of Kasumi-1 and NB4 compared to their shRecQ1 counterparts. Cells were untreated or treated for 6 h with 1µM olaparib. ≥100 fibers were scored per condition per replicate. Black line marks the median. Overlaid black dots represent the per-experiment medians (n = 3 for NB4; n=4 for Kasumi-1) ± SD. (C) Neutral comet OTM measurements for one biological replicate for Kasumi-1 and NB4 cells compared with their shRecQ1 counterparts. Cells were untreated or 1, 3 and 6 h after 1µM of olaparib treatment. ≥100 nuclei were scored per condition per replicate. Black line marks the median. Overlaid black dots represent the per-experiment medians (n = 3) ± SD. (D) Dose-response viability: NB4 and shRecQ1-NB4 treated with olaparib (1-15µM, 5 days). Cell numbers were quantified and normalized to the vehicle (DMSO) control. Dots and squares represent mean ± SD. N = 4 replicates. (B,C) Brown-Forsythe and Welch’s one-way ANOVA with Dunnett’s T3 multiple comparisons test. Indicated comparisons display numerical p-values.

These results indicate that PARPi sensitivity in defined AML subtypes is linked to deregulated RECQ1-mediated fork restart of reversed forks, leading to further fork acceleration and eventually to fork collapse.

### PrimPol levels determine replication fork integrity upon PARPi treatment in AML

As shown above, PARPi-resistant KMT2A-r AML cell lines (THP-1, MOLM-13) displayed no changes in fork speed or DNA damage levels upon olaparib treatment (Figure 1B, S1F). Hence, we set out to investigate how KMT2A-r AML cells would escape PARPi-mediated fork collapse. PrimPol-mediated repriming provides a mechanism alternative to fork reversal to deal with leading strand lesions (Bianchi et al., 2013; García-Gómez et al., 2013; Mourón et al., 2013). Because repriming competes with fork reversal and can enhance resistance to specific genotoxic treatments in solid tumours (Bai et al., 2020; Quinet et al., 2020), we reckoned that repriming may bypass reversal-based PARPi toxicity and thereby represent a PARPi resistance mechanism in KMT2A-r AML. RT-qPCR, western blotting, and analysis of public datasets revealed significantly higher PrimPol expression in KMT2A-r AML compared with RUNX1-RUNX1T1 and PML-RARɑ subtypes (Figure 3A and S2A). Moreover, proximity ligation assays (PLA) confirmed increased recruitment of PrimPol to EdU-labelled nascent DNA in KMT2A-MLLT3 relative to AE9a and PML-RARɑ mouse AML cells (Figure 3B).

**Figure 3:**
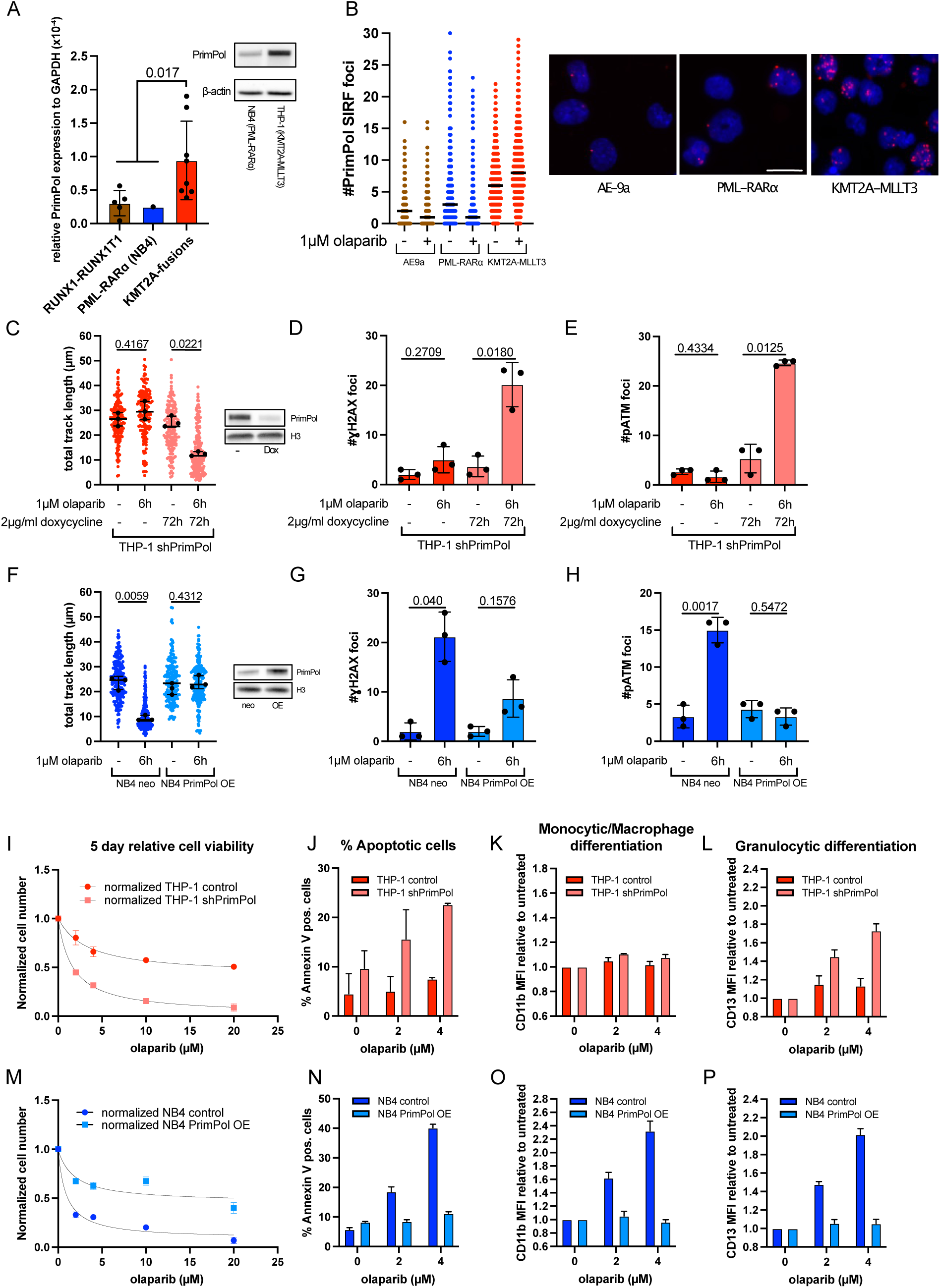
PrimPol levels determine PARPi molecular effects in AML. (A) RT-qPCR assay of *PRIMPOL* mRNA normalized to *GAPDH* in human AML cell lines and primary samples: KMT2A-rearranged (n=8) and CBF/APL (RUNX1-RUNX1T1 or PML-RARɑ; n=6) (left). PrimPol immunoblot comparing protein levels in NB4 and THP-1 (right). β -actin serves as loading control. (B) Left: Single replicate of in situ analysis of protein interactions at DNA replication forks (SIRF) foci counts quantifying PrimPol-EdU proximity in murine AML cell lines AE9a, PML-RARɑ and KMT2A-MLLT3 ± 1 µM olaparib (6 h). Right: representative images of PrimPol SIRF foci (red) over DAPI staining (blue). Scalebar, 10 µm. (C) THP-1 with doxycycline-inducible shPrimPol: Total fiber track lengths (left) for a representative biological replicate ± 1 µM olaparib (6 h) and ± doxycycline (72 h). ≥100 fibers were scored per condition per replicate. Black line marks the median. PrimPol immunoblot showing PrimPol depletion with H3 loading control (right). Overlaid black dots represent the per-experiment median from n = 3 independent experiments; error bars = SD of those medians. (D) ɣH2AX (Ser139) S-phase nuclear foci counts in THP-1 shPrimPol cells ± 1 µM olaparib (6 h) and ± doxycycline (72 h). ≥200 cells per condition per replicate. Black dots represent the per-experiment medians (n = 3). Bars represent mean ± SD. (E) pATM (S1981) S-phase nuclear foci counts in THP-1 shPrimPol cells, as in (D). ≥200 cells per condition per replicate. Black dots represent the per-experiment medians (n = 3). Bars represent mean ± SD. (F) NB4 ± PrimPol overexpression (PrimPol OE): Total fiber track lengths (left) for a representative biological replicate ± 1 µM olaparib (6 h). ≥100 fibers were scored per condition per replicate. Black line marks the median. PrimPol immunoblot confirming overexpression; H3 loading control (right). Overlaid black dots represent the per-experiment median from n = 3 independent experiments; error bars = SD of those medians. (G) ɣH2AX (Ser139) S-phase foci counts in NB4 and NB4 PrimPol OE ± 1 µM olaparib (6h). ≥200 cells per condition per replicate. Black dots represent the per-experiment medians (n = 3). Bars represent mean ± SD. (H) pATM (S1981) S-phase foci counts in NB4 and NB4 PrimPol OE, as in (G). ≥200 per condition per replicate. Black dots represent the per-experiment medians (n = 3). Bars represent mean ± SD. (I) Cell-count dose-response curves for olaparib (0-20µM, 5 days) in THP-1 WT and THP-1 doxycycline-inducible shPrimPol (THP-1 shPrimPol) cells. (J) Annexin V-positive fraction after 5 days of olaparib (0-4µM) in THP-1 WT and THP-1 shPrimPol. (K) CD11b mean fluorescence intensity (MFI), relative to untreated, after 5 days of olaparib (0-4µM) in THP-1 WT and THP-1 shPrimPol. (L) CD13 MFI, relative to untreated, after 5 days of olaparib (0-4µM) in THP-1 WT and THP-1 shPrimPol. (M) Cell-count dose-response curves for olaparib (0-20µM, 5 days) in NB4 WT and NB4 PrimPol overexpressing (PrimPol OE) cells. (N) Annexin V-positive fraction after 5 days of olaparib (0-4µM) in NB4 WT and NB4 PrimPol OE. (O) CD11b mean fluorescence intensity (MFI), relative to untreated, after 5 days of olaparib (0-4µM) in NB4 WT and NB4 PrimPol OE. (P) CD13 MFI, relative to untreated, after 5 days of olaparib (0-4µM) in NB4 WT and NB4 PrimPol OE. (A) Unpaired Welch’s t-test. (C-H) Brown-Forsythe and Welch’s one-way ANOVA with Dunnett’s T3 multiple comparisons test. Indicated comparisons display numerical p-values. (I-P) Unless indicated otherwise, bars and dose-response curve symbols indicate mean ± SD of n = 2 biological replicates.

To test whether elevated PrimPol activity in KMT2A-r AML underlies PARPi resistance, we conditionally depleted PrimPol in KMT2A-MLLT3 cells. PrimPol knockdown combined with olaparib treatment in both THP-1 and MOLM-13 caused pronounced shortening of replicated tracks, accompanied by increased S-phase DNA damage and DSB signalling (Figure 3C-E and S2B-D). Conversely, constitutive PrimPol overexpression in PARPi-sensitive cell lines NB4 and Kasumi-1 rescued replication tract lengths, markedly reduced S-phase DNA damage signalling and abolished DSB signalling upon olaparib treatment (Figure 3F-H, S2E-G).

These findings identify PrimPol as a key modulator of PARPi response in AML and suggest that elevated PrimPol activity contributes to PARPi resistance in KMT2A-r subtypes.

### PrimPol promotes resistance to PARP inhibitors in *in vitro* AML models

We next asked whether the striking mechanistic differences in replication dynamics and DNA damage accumulation among the AML cell lines analyzed – largely reflecting a different balance between fork reversal and repriming – would impact long-term survival and differentiation upon extended PARPi treatment. In line with our short-term mechanistic findings, THP-1 cells displayed reduced olaparib sensitivity, which could be markedly increased by PrimPol depletion (Figure 3I), associated with increased apoptosis and granulocytic differentiation (Figure 3J-L). Similarly, although to a minor extent, PrimPol depletion sensitized MOLM-13 cells to olaparib exposure (Figure S2H). Conversely, prolonged olaparib exposure in NB4 and Kasumi-1 cells significantly reduced viability, accompanied by increased apoptosis and induction of monocytic/macrophage and granulocytic differentiation. Strikingly, PrimPol overexpression in both AML lines largely rescued cell viability and impaired apoptosis/differentiation under these conditions (Figure 3M-P, S2I-L).

### PrimPol sustains fork progression and cellular resistance upon araC treatment

PrimPol can reprime downstream of a wide range of replication-blocking lesions and obstacles (Guilliam & Doherty, 2017; Tirman, Cybulla, et al., 2021). Notably, PrimPol has been shown to bypass incorporated chain-terminating nucleoside analogues, allowing replication to continue despite the presence of such roadblocks (Kobayashi et al., 2016). Given that Cytarabine (araC) represents a standard chain-terminating nucleoside analogue used in AML therapy, we asked whether PrimPol activity might also modulate the cellular responses to this drug. As expected, short (6 h) low-dose (10 nM) araC treatment significantly shortened replicated tracks in both NB4 and Kasumi-1 cells. Remarkably, replication fork slowdown in both cell lines was fully rescued by PrimPol overexpression (Figure S3A,B).

Immunofluorescence analysis of NB4 cells revealed reduced DNA synthesis (EdU levels), and increased DNA damage signalling (ɣH2AX foci) following araC treatment, but no detectable DNA break accumulation (pATM; Fig. S3C-F). PrimPol overexpression rescues DNA synthesis in the presence of araC (Fig. 3D) and significantly mitigates araC sensitivity in both NB4 and Kasumi-1 cells, by drastically reducing drug-induced differentiation, while apoptosis is not significantly affected (Fig. S3G-N).

Hence, PrimPol facilitates the bypass of chain-terminating nucleoside analogues such as araC, contributing to resistance of defined AML lines against standard-of-care AML therapies.

### High PrimPol expression confers chemoresistance in AML

These promising observations on AML cellular models prompted us to assess whether PrimPol could be considered a *bona fide* a drug-resistance factor in AML. TARGET-AML survival analysis of primary peripheral-blood samples (all ages) revealed that higher PrimPol expression is indeed associated with significantly worse overall patient survival, likely reflecting limited or less durable therapy response (Figure 4A). We next tested in xenotransplantation mouse model whether PrimPol expression also impacts treatment outcome *in vivo*. NB4 cells with and without PrimPol overexpression were transplanted into immunodeficient NBSGW mice, which were subsequently treated with vehicle, olaparib or araC (Figure 4B). Mice transplanted with wild-type NB4 cells survived significantly longer following araC or olaparib treatment (Figure 4C). Strikingly, overexpression of PrimPol abolished these survival benefits, with no significant improvement observed upon either treatment (Figure 4D).

**Figure 4:**
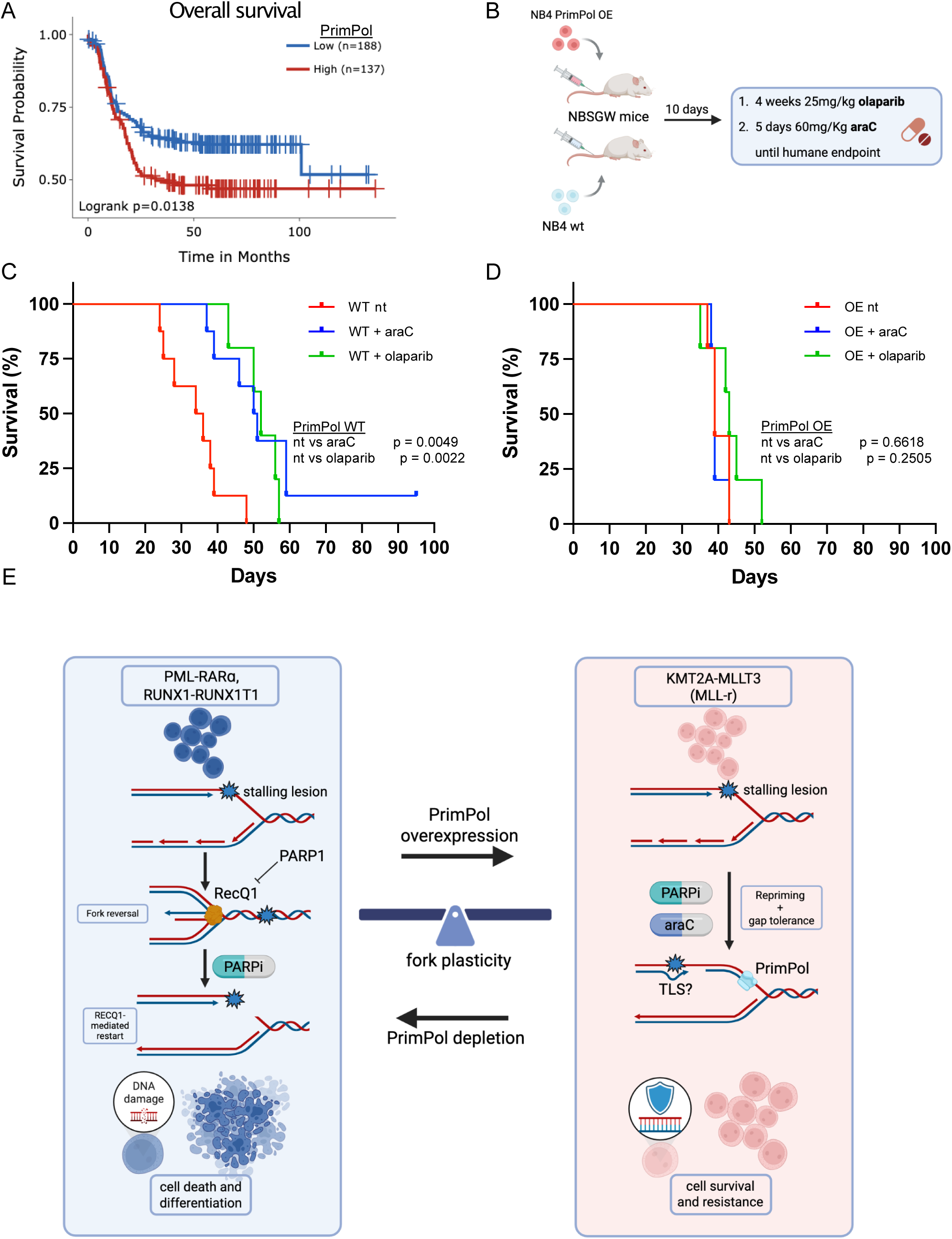
High PrimPol expression counteracts therapy response in AML. (A) Kaplan-Meier overall survival in TARGET-AML (primary peripheral-blood samples; all ages), stratified by PRIMPOL expression using the single-gene module with optimal cut-point determination (Survival Genie v2.0). Curves compare PRIMPOL-high vs. PRIMPOL-low; log-rank P = 0.0138; n (high/low) = [137/188]. (B) NBSGW mouse experimental scheme. Mice were transplanted with NB4 WT or NB4 PrimPol overexpressing cells. Ten days post-transplant, cohorts received vehicle, olaparib 25mg/kg for 4 weeks, or cytarabine 60mg/kg for 5 days, and were monitored to humane endpoint. (C) Kaplan-Meier survival curves for NBSGW mice bearing NB4 WT cells under the treatments in (B); n = 8 mice (vehicle, araC) and n = 5 mice (olaparib) per group. (D) Kaplan-Meier survival curves for NBSGW mice bearing NB4 PrimPol overexpressing cells under the treatments in (B); n = 5 mice per group. (E) Fork plasticity balance model in AML. Replication forks in AML balance fork reversal/restart (RECQ1-dependent) against PrimPol-mediated repriming. In reversal-leaning AML subtypes (e.g. RUNX1-RUNX1T1, PML-RARɑ) PARP1 PARylation stabilizes reversed forks by limiting RECQ1 restart. PARP inhibition (PARPi) forces fork restart and acceleration, culminating in fork collapse and DNA damage accumulation. In repriming-leaning AML (e.g., KMT2A-MLLT3), forks favor repriming, deferring lesions to post-replicative repair gap filling, which mitigates PARPi-induced cytotoxicity. This framework explains the divergent olaparib (and cytarabine) responses observed across cell lines. (C,D) Differences between Kaplan–Meier survival curves were assessed using the Logrank (Mantel–Cox) test, with two-sided *p* values < 0.05 considered statistically significant.

Together, these findings establish PrimPol as a critical determinant of chemotherapeutic resistance and differentiation blockade in AML.

## Discussion

Our dataset establishes a novel link between drug response in AML and differential usage of DNA damage tolerance mechanisms at replication forks. The endogenous lesions inducing frequent fork reversal or PrimPol-mediated repriming in hyper-proliferating myeloid precursor cells are currently unclear. Reactive oxygen species, endogenous aldehydes (formaldehyde) and transcription-replication conflicts were all suggested to represent a challenge for DNA replication and genome stability in proliferating HSPCs (Chen et al., 2020; Ito et al., 2006; Pontel et al., 2015; Tothova et al., 2007; Wang et al., 2022). Assessing their relative abundance and mechanistic contribution upon the different epigenetic deregulations driving AML will provide important molecular clues to understand the replication fork transactions observed in each AML subtype and to better exploit them for therapeutic purposes.

In t(8;21) AML (RUNX1-RUNX1T1) and APL (PML-RARɑ), replication forks are apparently prone to undergo frequent fork reversal, exposing them to PARPi-mediated deregulation of RECQ1-dependent fork restart. Conversely, KMT2A-r AML cells avoid fork reversal and extensively engage PrimPol in DNA synthesis; this discontinuous replication mode protects these AML cells from PARPi effects on reversed fork restart and assists lesion bypass also in response to standard-of-care genotoxic treatments, helping to explain the reported chemoresistance of KMT2A-r (van Weelderen et al., 2024; Zeisig et al., 2012); Figure 4E). Based on the mechanistic knowledge available on alternative mechanisms of reversed fork processing and restart, it seems likely that, besides RECQ1, other fork processing mechanisms – such as nuclease-mediated cleavage (MUS81-EME2) and/or MRE11/DNA2 processing of regressed arms (Lemaçon et al., 2017; Pepe & West, 2014; Regairaz et al., 2011; Thangavel et al., 2015) – play important roles in the observed DSBs and cytotoxicity induced by PARPi or standard-of-care AML treatments. Besides the effects on reversed fork restart, PARPi may be effective in the sensitive cell lines by increasing the collisions of accelerated forks with trapped PARP1/2 protein-DNA roadblocks and/or with other replication/repair intermediates known to accumulate upon PARPi treatment, such as unligated Okazaki fragments, TOP1 cleavage complexes, BER intermediates or transcription-replication conflicts (Berti et al., 2013; Chowdhuri & Das, 2021; Murai et al., 2012; Vaitsiankova et al., 2022). Moreover, fork acceleration upon defective PARylation limits checkpoint signalling, which may render replication forks intrinsically more unstable (Calzetta et al., 2025; Maya-Mendoza et al., 2018). Further investigation is needed to assess the relative contribution of these and other mechanisms in the chemosensitivity of these AML subtypes and will likely help identifying new actionable entry points for therapeutic intervention.

By contrast, KMT2A-MLLT3 (t(9;11)) – and possibly other KMT2A-r AMLs – favour PrimPol-mediated repriming under replication stress, enabling rapid traverse of blocking lesions at the cost of post-replicative ssDNA gap accumulation (Figure 6E). In solid tumours, ssDNA gaps have recently emerged as key lesions underlying PARP-inhibitor cytotoxicity, providing a therapeutic rationale for inducing more gaps and/or preventing their repair (Cong & Cantor, 2022; Quinet et al., 2021; Vaitsiankova et al., 2022). Our findings in AML are particularly surprising in this respect, since KMT2A-MLLT3 models extensively use PrimPol-dependent discontinuous synthesis and are yet relatively insensitive to PARP inhibition. To reconcile these observations, we propose that AML subtypes engaging PrimPol in DNA synthesis are likely very efficient in filling the transient ssDNA gaps resulting from PrimPol function. Several mechanisms are known to mediate ssDNA gap repair, i.e. translesion synthesis (REV1/Polζ), Polθ-mediated end joining, and/or RAD52-dependent DNA synthesis operating after bulk DNA replication (MiDAS/BIR-like repair) (Bhowmick et al., 2016; Schrempf et al., 2022; Sotiriou et al., 2016; Tirman, Quinet, et al., 2021). Among these, Polθ is a compelling candidate mechanism mediating chemoresistance in AML, as it is frequently upregulated across AML subsets and it has shown promise in preclinical studies on AML (Sullivan-Reed et al., 2023; Vekariya et al., 2023). Polθ operates independently of PCNA ubiquitination, by direct recruitment to RPA- or RAD51-coated ssDNA, catalysing microhomology-mediated annealing and 3’ extension (Schaub et al., 2022; Vekariya et al., 2023). Based on our evidence on PrimPol engagement in specific AML subtypes, assessing ssDNA gap persistence upon Polθ inactivation may illuminate novel strategies to tackle the reported chemoresistance of KMT2A-r AML.

Our data identify RECQ1 and PrimPol – and particularly their chromatin loading or binding to replication forks – as potential predictive biomarkers for therapy response and acquired chemoresistance in AML. High RECQ1/low PrimPol engagement – as for t(8;21) and APL – would identify AML relying on the reversal-RECQ1-restart axis, likely to benefit from PARPi-mediated therapies. In these AMLs, combining PARP inhibitors with classical chemotherapeutics inducing DNA adducts or topoisomerase lesions (e.g. camptothecin or decitabine) is likely to enhance the effects of PARPi-driven fork restart and fork breakage, contributing to potent cytotoxicity. Conversely, high chromatin/fork binding of PrimPol would predict chemoresistance, particularly if accompanied by a high “gap filling” signature (Polθ, TLS), suggesting alternative therapeutic strategies that target these enzymes. In these cases, classical replication-blocking chemotherapies should be combined with gap-filling inhibitors to increase fork roadblocks and ssDNA gap load, while preventing their repair (Hollenbach et al., 2010; Patel et al., 2012). Direct PrimPol inhibitors are unfortunately not yet available, and our work clearly suggests that this should be an important strategy to explore for AML therapy. However, until such a specific inhibitor is available, the use of ATR or CHK1 inhibitors in combination with PARPi may prevent PrimPol recruitment as well as proper ssDNA gap repair, thereby increasing fork collapse and drug cytotoxicity (Mehta et al., 2022; Wethington et al., 2023). Moreover, parallel work within our team has recently identified DOT1L/menin inhibitors as effective epigenetic regulators counteracting Primpol-dependent repriming in favour of replication fork reversal, offering a valuable and clinically attractive strategy to sensitize chemoresistant KMT2A-r AML to PARPi treatment both *in vitro* and *in vivo* (ECW So, unpublished observations). Overall, these mechanistic studies pave the way for novel therapeutic strategies in AML, likely to improve patient stratification and clinical outcome, based on the molecular profiling of replication fork plasticity.

## Acknowledgements

We thank all members of the Lopes laboratory for scientific input, helpful discussions and general support throughout this project. This work was supported by the Swiss National Science Foundation (SNSF) Sinergia grant (CRSII5_213586 / 1).

## Author Contributions

**C.D.** in part conceived the mechanistic study framework, designed, performed, and analyzed all mechanistic experiments, and wrote the manuscript. **T.K.F.** performed most in vitro and in vivo treatment sensitivity assays and contributed to cell line generation. **T.G.** performed experiments and provided troubleshooting support. **L.M.B.** designed and performed in vivo experiments and analyzed in vivo data. **C.W.E.S.** contributed to funding, study design and supervision, and manuscript editing. **M.L.** conceived and supervised the project, secured funding, and contributed to study design, data interpretation, and manuscript writing/editing.

## Methods

### Retroviral transduction and transformation assay (RTTA)

To generate murine AML primary cell lines, RTTA was applied to express oncogenes (Ae9a, PML-RARɑ or KMT2A-MLLT3) in mouse HSPCs. Mouse bone marrow (BM) was harvested from femur and tibia of wild-type C57bl/6 mice. BM was dissociated in 10 mL PBS + 2% FBS and spun down, followed by red cell lysis to remove red blood cells by incubate BM cells in 1mL lysis buffer for 5 mins. Cells were then washed again in 10 mL PBS+2% FBS and resuspended in 0.5 mL. To isolate HSPCs (c-Kit+ cells) magnetic activated cell sorting (MACS) was used, incubating BM cells with 20µl of anti-CD117 magnetic microbeads (Miltenyi Biotec; 130-091-224) for 20 min in the fridge. Cells were then washed with 10 mL PBS+2% FBS and resuspended in 1 mL. After washing, cells were applied to MS MACS column (Miltenyi Biotec; 130-042-201) for MACS according to manufacturer instruction to purify c-Kit+ cells.

Twenty-thousand c-Kit+ cells were subsequently seeded in U-shaped 96-well plates and cultured overnight in RPMI medium supplemented with 10ng/mL mouse IL-3, IL-6 and SCF. The following day, cells were incubated with 200µl retroviral particles expressing the oncogenes, supplemented with 10ng/mL mouse IL-3, IL-6 and SCF, and then spinoculated for 2 hours at 32oC at 800 x g. After spinoculation, cells were placed into CO2 incubator overnight.

The next day transduced cells were plated into methocult medium (StemCell Biotechnology; M3434) with 10ng /mL of mouse IL-3, IL-6, SCF and GM-CSF, with the addition of 1.5ug/mL puromycin to select transduced cells. After 7-10 days, cell colonies start to appear in methocult medium. Subsequently, colonies are harvested and washed in 10 mL PBS+2% FBS. 5K of MLLT3-KMT2A or 20K of AE9a or PML-RARɑ transformed cells, were seeded into methocult as above for a 2nd replating and repeated again for an additional 3rd replating. HSPCs were considered transformed into AML primary cell lines after 3rd replating.

### Cell culture

Human NB4, Kasumi-1, THP-1 and MOLM-13 cell lines (all from DMSZ) were cultured in RPMI 1640 (Gibco, 11875093) with the addition of 10% fetal bovine serum (FBS), 1% penicillin/streptomycin (Sigma-Aldrich, 12352207) and 10ng/mL each of human IL-3 (Peprotech, 200-03), IL-6 (Peprotech, 200-06), SCF (Peprotech, 300-07), FLT3 (Peprotech, 300-19) ligand and TPO (Peprotech, 300-18). Cells were cultured at 5% CO2 and 37°C. Healthy donor peripheral blood and bone marrow samples were collected from blood donors by the Blutspende Zürich under a study protocol approved by the cantonal ethics committee, Zürich (KEK Zürich, BASEC-Nr 2019-01579). Human healthy donor CD34+ samples were thawed in thawing media containing 12.25ml RPMI medium + 12.25ml FBS + 1ml 5mg/ml DNAse 1 solution (Merck, 11284932001) and after centrifugation at 200g for 10min, cells were resuspended in culture media containing 2ml X-Vivo10 media (Lonza, BP04-743Q) per well, supplemented with 20µl 100X GlutaMAX (Gibco, 35050-061)+ 10µg/ml SCF + 50µg/ml TPO + 10µg/ml Flt3. Cells were cultured for 48h before use. RTTA-obtained mouse MLL-AF9 and Ae9a cells were cultured in RPMI 1640 (Gibco, 11875093) with the addition of 20% fetal bovine serum (FBS), 20% WEHI-conditioned medium (generated by So lab) and 1% penicillin/streptomycin (Sigma-Aldrich, 12352207). Mouse PML-RARɑ was cultured in IMDM (Thermo Fisher Scientific, 31980030) with 15% FBS and 1% penicillin/streptomycin. Additionally, the mouse AML media was supplemented with 10ng/mL each of mouse IL-3 (Peprotech, 213-13), IL-6 (Peprotech, 216-16) and SCF (Peprotech, 250-03).

### Drugs and ionizing irradiation

Olaparib was ordered from LubioScience (T3015) or Selleckchem (S1060), Cytarabine (S1648) was ordered from Selleckchem.

For positive DNA damage control, AML cells were irradiated with x-rays using a RadSource 2000 (LASC UZH Irchel) for 5-10 Grey irradiation as indicated per experiment.

### Plasmid transfections

PrimPol shRNA plasmid was constructed in the lab of Juan Méndez as part of (Mourón et al., 2013). THP-1 and MOLM-13 cells were transduced in the So lab with a TRIPZ lentiviral vector carrying an inducible shRNAmir targeting the sequence 5’ TGCTGTTGACAGTGAGCG in the 3’ untranslated region (UTR) of the PRIMPOL gene (Open Biosystems). Transduced cells were selected using 1.5ug/mL puromycin (ant-pr-1). Expression of PRIMPOL shRNAmir was induced with 1 μg/mL doxycycline (Dox) (Merck, D9891) for 3 days. Cells expressing PRIMPOL shRNA are RFP+.

NB4 and Kasumi-1 cells with stable PRIMPOL overexpression were generated by retroviral transduction in the So lab, using MSCV plasmid containing a neomycin resistance and expressing human PRIMPOL cDNA. Transduced cells were selected using 1mg/mL G-418.

RECQL shRNA plasmid was constructed targeting the RECQL sequence 5’-CCG GGC ACA TGC TAT TAC TAT GCA ACT CGA GTT GCA TAG TAA TAG CAT GTG CTT TTT G-3’ (TRCN0000289591). Human RECQL shRNA was cloned into a pLKO.1 plasmid. Lentivirus was produced by transfecting HEK 293T cells with shRNA containing pLKO.1, pVSV, pMDL and pREV lentiviral packaging plasmids. After lentivirus collection NB4 and Kasumi-1 cells were transduced and selected using puromycin (ant-pr-1). The depletion of RECQ1 was tested using immunoblotting.

### DNA fiber assay

Human or mouse AML cells were optionally treated with drugs at indicated doses (see figure legends) prior to labelling. Human AML cell lines and healthy donor CD34+ cells were pulse-labelled in their respective culture media with 0.045mM 5-Chloro-2’-deoxyuridine (CldU; Sigma-Aldrich, C6891) for 30 min. After centrifugation at 400g, media containing 0.31mM 5-iodo-2’-deoxyuridine (IdU; Sigma-Aldrich, 17125) was added to cells for 30 min. Due to IdU toxicity nucleotide concentrations were adjusted in mouse leukemia cell lines to 0.026mM CldU and 0.186mM IdU, with centrifugation speeds not exceeding 200g. Subsequently, DNA fibers were spread as previously described (Jacobs et al., 2022). In brief, 3µl cell suspension (1 million cells/mL) was added to 7µl lysis buffer (50mM EDTA, 0.5% (W/V) SDS and 200mM Tris-HCl, pH=7.5) on glass slides. After 5 min of lysis, fibers were spread by tilting the slides at an angle (30-45°). After air-drying the slides were fixed in -20°C 3:1 methanol/acetic acid for 10 min. The fibers were then denatured with 2.5M HCl for 1h, washed with PBS to neutralize the pH and blocked for 45 min with blocking solution (PBS 1X, 0.2% Tween20 (PBS-T), 1% BSA (Merck, 05470)). Slides were incubated with anti-BrdU antibodies binding IdU (BD Biosciences, 347580, 1:100) and CldU (Abcam, ab6326, 1:300) in the dark for 3h at room temperature (RT), followed by washing with PBS-T and 1.5h incubation with secondary antibodies at RT in the dark: goat anti-rat Alexa Fluor 555 (Thermo Fisher Scientific, A-21434, 1:300) and goat anti-mouse Alexa Fluor 488 (Thermo Fisher Scientific, A-11001, 1:300). After washing with PBS-T, slides were mounted with Prolong Gold antifade (Thermo Fisher Scientific, P36930) and imaged using a Leica DM6 microscope (HCX PL APO, 63x, oil). Fiber images were analysed using ImageJ software.

### Immunofluorescence staining of ɣH2AX and pATM

After varying treatments, cells were incubated with 25µM 5-ethynyl-2’-deoxyuridine (EdU; Thermo Fisher Scientific, A10044) in the last 10 min before collection. Cells were optionally irradiated with x-rays using a RadSource 2000 for 10 Grey irradiation after a 10 min 25µM EdU pulse. Cells were subsequently sedimented on poly-L-lysine (Merck, P4832) pre-coated VWR 13mm coverslips, placed inside 24-well plates. To sediment cells, the coverslip containing plates were spun at 400g for 5 min. Cells were pre-fixated in 0.1% formaldehyde (Sigma-Aldrich, F8775), pre-extracted using ice-cold CSK-buffer (10mM 4-(2-hydroxyethyl)-1-piperazineethanesulfonic acid (HEPES) pH=7.4, 0.1M NaCl, 0.3M sucrose, 3mM MgCl2, 0.5% Triton-X (Sigma-Aldrich, 93443)) for 5 min and fixed for 15min using 4% formaldehyde (Sigma-Aldrich, F8775). Cells were then blocked using 5% BSA (Merck, 05470) + 10% fetal bovine solution (FBS) for 1h at 37°C and incubated with rabbit anti-ɣH2AX S139 (cell signalling technology, 9718S, 1:200) and mouse anti-pATM S1981 (Rockland Immunochemicals, 200-301-400, 1:1000) primary antibodies overnight at 4°C. Subsequently, cells were washed with PBS and incubated for 1h at 37°C with blocking solution containing Alexa Fluor 488 anti-mouse (Thermo Fisher Scientific, A11001) and 647 anti-rabbit (Thermo Fisher Scientific, A31573) secondary antibodies at 1:300 dilution. After PBS washes, EdU containing cells were stained for 30 min using 100µl homemade Click-iT Cell reaction buffer (100mM Tris pH=8, CuSO4, 10 mM sodium-L-ascorbate) with 0.5µl Alexa Fluor 555 azide (Thermo Fisher Scientific, A20012) per coverslip. After 1µg/ml 4’, 6-diamidino-2-phenylindole (DAPI; Merck, D9542) staining, washing and mounting with Prolong gold antifade (Thermo Fisher Scientific, P36930), coverslips were imaged using a Leica DM6 microscope (HCX PL APO, 63x, oil) and analyzed using Olympus ScanR Image Analysis software (Version 3.3.0, 3.0.1). Using the ScanR software, edge-based background correction for nuclei and foci detection masks was applied equally to all compared conditions per replicate. Foci counts and fluorescence intensities were quantified and exported to be analyzed using TIBCO Spotfire software (Version 10.10.1).

### Neutral comet assay

After treatment, 1000-2000 cells per condition were embedded in 37°C 0.8% SeaPlaque low-melting point agarose (Lonza, 50111) per gel on 2-well comet slides (Trevigen, 4250-200-03). Cells were lysed overnight at 4°C in lysis buffer (2.5M NaCl, 100mM EDTA and 10mM Tris (pH=10)), supplemented with 10% DMSO and 1% Triton X-100 (Sigma-Aldrich, 93443). Cells were optionally irradiated with x-rays using a RadSource 2000 for 10 Grey irradiation before embedding. For neutral comet assays, the slides were washed and incubated for 1h in neutral electrophoresis buffer (300mM Sodium Acetate, 100mM Tris, pH=8.3) followed by electrophoresis for 35 min at 25V and 300mA using a CometAssay electrophoresis system (Bio-techne, 4250-050-ES). The slides were then washed with PBS and then immersed in DNA precipitation solution (1M NH4Ac, 95% Ethanol) at room temperature (RT) for 30 min. Slides were then placed into 70% ethanol for 30 min at RT and subsequently dried at 37°C. DNA was stained with SYBR gold (Thermo Fisher Scientific, S11494) for 30min, followed by PBS wash and drying at 37°C. The slides were imaged using an IN Cell Analyzer 2500 HS (GE Healthcare Life Sciences) and scored using the Open Comet plugin available for Fiji (ImageJ).

### Cell sensitivity assay

Cells were seeded in 96 – or 48 – well plates with a cell density of 100k-200k/mL in culture medium and treated with drugs at the doses indicated in figures. After drug incubation, cells were transferred into Eppendorf tubes, washed and resuspended thoroughly by vortexing. 10 µl of cell suspension was mixed with 10 µl of Trypan Blue solution. 10 µl of the mixture was then applied to a hematocytometer and counted under a cell culture microscope. Cell growth upon drug treatment was normalized to control DMSO treated cells. Assays were performed in duplicates.

### Annexin V apoptosis assay

Cells were harvested into Eppendorf tubes and washed. Cells were then resuspended in 300 µl of 1X Annexin V buffer (Thermo Fisher; 00-0055-56) and 100 µl of resuspended cells were transferred to FACS tubes. 100 µl of antibody mix, containing 1:100 APC-Annexin V antibody (Biolegend; 640920), and 1:1000 PI in PBS, was added and the cells were incubated at room temperature for 30 mins. Cells were subsequently washed with 2 mL 1X Annexin V buffer and resuspended in 400 µl 1X Annexin V buffer. Stained cells were then subjected to flow cytometric analysis using BD LSRII, using mean fluorescence intensity. Apoptotic cells were scored as APC-positive in histogram. Changes in marker expression upon drug treatments were normalized to that of DMSO control treated cells.

### FACS assay of differentiation

Cells were harvested into Eppendorf tubes and washed with PBS after treatment with drugs at the doses indicated in figures. Cells were resuspended in 300 µl of PBS and 100 µl of resuspended cells were transferred into FACS tube. 100 µl of antibody mix, containing 1:100 APC-CD13 (Biolegend;301706) and PE-Cy7-CD11b (Biolegend; 101216) in PBS, was added to the cells and incubated in the fridge for 30 mins. Cells were subsequently washed with 2 mL PBS, and resuspended in 400 µl PBS. Stained cells were subjected to flow cytometric analysis using BD LSRII. Expression of cell surface markers were analysed using mean fluorescence intensity. Change in marker expression upon drug treatments were normalized to that of DMSO control treated cells.

### Reverse transcription quantitative polymerase chain reaction (RT-qPCR)

RNA was isolated from cell lines or primary patient samples using a total RNA extraction kit (NEB; T2010S) according to the manufacturers instructions. 0.5 to 1µg of total isolated RNA were subjected to reverse-transcription reaction using SuperScript™ III Reverse Transcriptase (Invitrogen; 18080044) with oligo-dT (18mers) according to the manufacturers instructions. cDNAs were then used to perform quantitative PCR (qPCR) using Fast SYBR™ Green Master Mix (Thermo Fisher; 4385614) according to the manufacturers instructions, with a StepOnePlus System (Applied Biosystems) by in-build FAST running cycle. GAPDH was used as the house-keeping gene to normalize the expression of PRIMPOL, and Primpol expression was presented as log2 (deltaCt of GAPDH – PRIMPOL). Primer sequences as below:

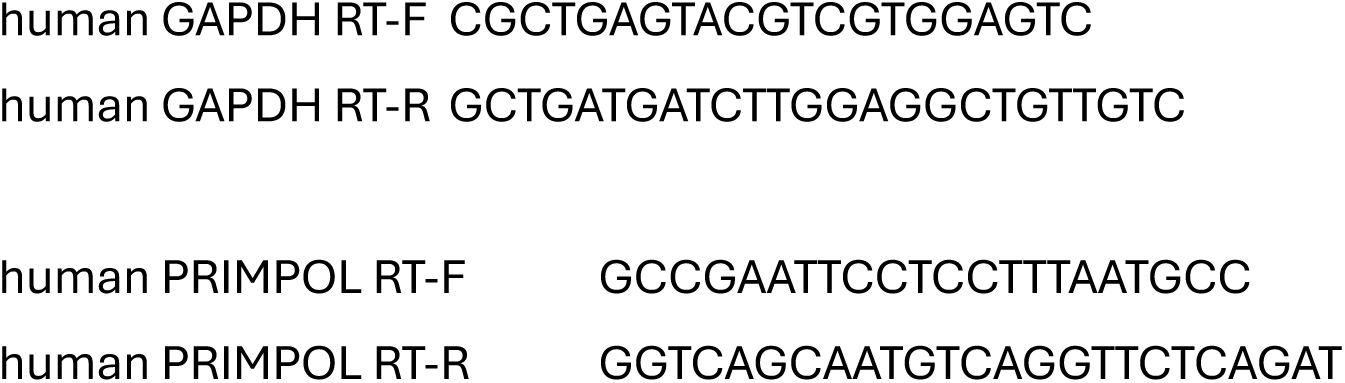

### Murine in vivo sensitivity experiments

200K wild-type NB4 cells or NB4 cells with PRIMPOL overexpression were resuspended in 150 µl PBS, then transplanted via tail vein into immunodeficient NBSGW mice (Jax ID: 026622). 10 days after transplantation, mice were either mock treated, or treated with 25mg/Kg olaparib (in 10% w/v (2-Hydroxypropyl)-β-cyclodextrin (HBC) in water; Merck; H107-5G) for 4 weeks or 60mg/Kg araC (in PBS) for 5 days. After drug treatment, mice were monitored for any sign of sickness, e.g. weight loss, hunch posture, slow movement, dragging hind leg(s), any tumour lump formed. Mice were culled once close to humane endpoint as protocol allows. Animals were considered AML bearing, if they were showing any signs of sickness mentioned above. This include anatomy of enlarged spleen and/or liver, and/or tumour formed, with infiltration of human AML cells into any tissues/ tumours, evaluated by FACS.

BM, spleen, liver, blood and tumour (if formed) were harvested for FACS analysis to detect human AML cells. Human engraftment is determined by FACS with human CD45+ human CD33+.

### TARGET AML dataset survival analysis (Survival Genie v2.0)

Patient data Kaplan-Meier survival curves were generated with the single-gene analysis module of Survival Genie v2.0 (Bhasin lab, Emory university). Analysis was restricted to the TARGET-AML cohort, primary samples from peripheral blood, all age groups. For PrimPol, patient groups were define using the platform’s optimal cut-point determination (maximally selected rank statistic), and survival differences were evaluated by a two-sided log-rank test. The plot was exported directly from the web interface (April, 2025) and used without further graphical modification.

### Statistical analyses

Statistical analysis was performed as mentioned in the figure legends, using GraphPad Prism software (version 10.4.2)

All experiments were statistically evaluated using either unpaired Welch’s t-test, Brown-Forsythe and Welch’s one-way ANOVA with Dunnett T3 multiple comparisons, or Logrank (Mantel–Cox) test, as indicated in the figure legends. P<0.05 was deemed statistically significant.

