## Supplementary Figures 1-3 for "Replication fork plasticity is a therapeutic vulnerability in acute myeloid leukemia"

### Supplementary Figures and figure legends

Figure S1:

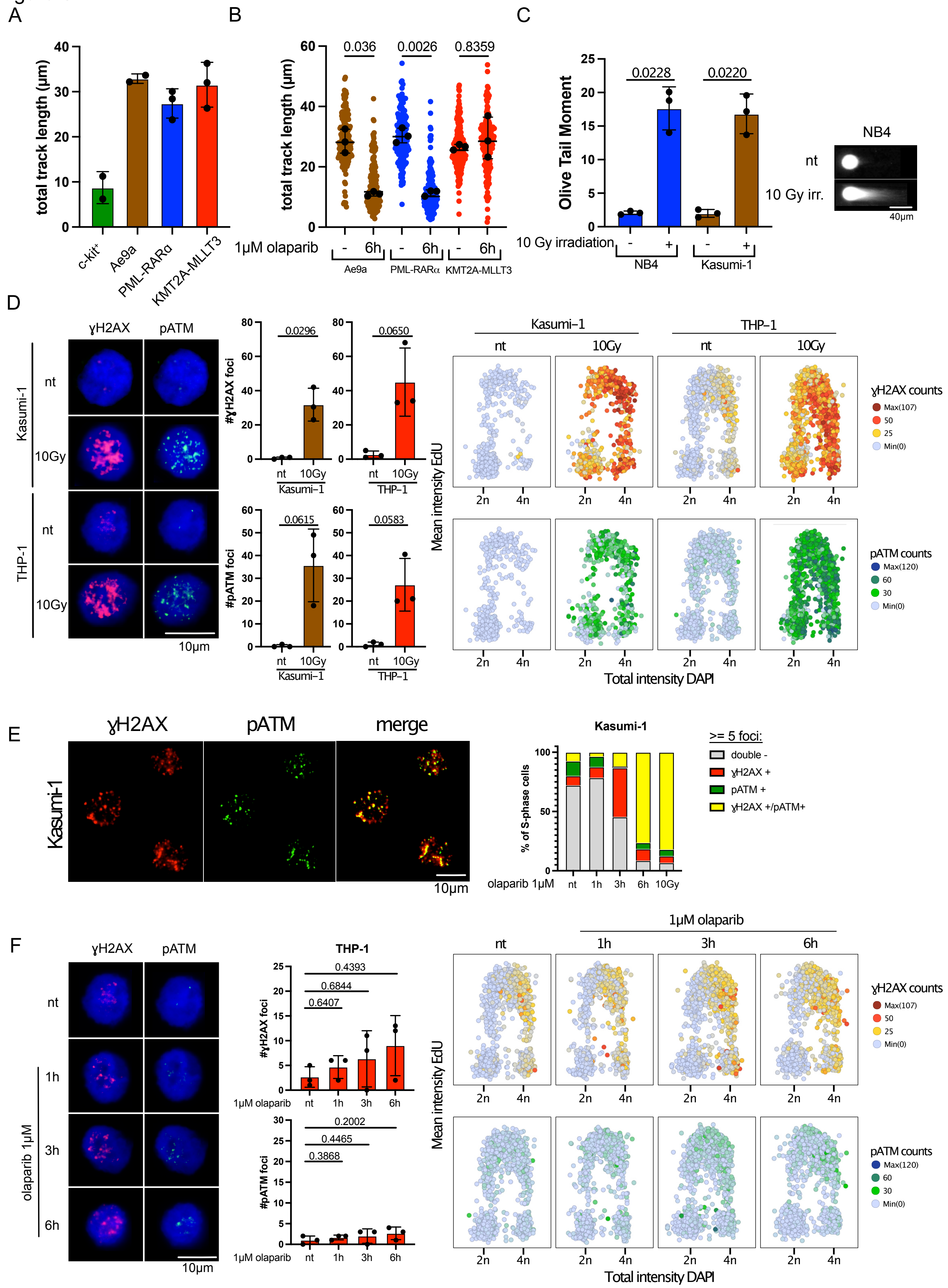

**Figure S1** (related to Figure 1).

(A) Total CldU + IdU fiber track lengths in c-kit$^+$ murine hematopoietic cells and retrovirally transduced/transformed mouse AML cell lines (Ae9a [RUNX1-RUNX1T1(9a)], PML-RARα, KMT2A-MLLT3). ≥100 fibers were scored per condition per replicate. Black dots represent the per-experiment median from n = 2 (c-kit$^+$, Ae9a) and n= 3 (PML-RARα, KMT2A-MLLT3) independent experiments; Bars represent mean ± SD of those medians.

(B) Total fiber track lengths for a representative biological replicate of murine AML cell lines, untreated (-) or treated with 1μM olaparib for 6h. Black line marks the median. Overlaid black dots represent the per-experiment median from n = 3 independent experiments; error bars = SD of those medians.

(C) Neutral comet assay irradiation positive control: representative comets (right) and Olive Tail Moment (OTM) quantification (left) in Kasumi-1 and NB4 upon 10 Gy ionizing radiation. ≥100 nuclei were scored per condition per replicate. Black dots represent the per-experiment median from n = 3 independent experiments. Bars represent mean ± SD. Scalebar, 40 μm

(D) Positive-control IF/QIBC: THP-1 and Kasumi-1 untreated or after 10 Gy IR. Representative images (left), S-phase nuclear foci counts for ɣH2AX and pATM (middle) and QIBC DNA-content (total DAPI intensity) vs. DNA-synthesis (log-mean EdU intensity) profiles (right). Per-cell ɣH2AX (top) and pATM (bottom) foci counts are encoded by color. ≥ 200 cells were quantified per condition per replicate. Black dots represent the per-experiment medians (n = 3). Bars represent mean ± SD. Scalebar, 10 μm.

(E) Left: representative IF images of ɣH2AX (Ser139) and pATM (S1981) in S-phase Kasumi-1 cells after 6 h of 1μM olaparib. Yellow color in the merged image indicates colocalization of ɣH2AX and pATM foci. Scalebar, 10 μm. Right: quantification of one representative biological IF replicate of S-phase Kasumi-1 cells co-stained for ɣH2AX (Ser139) and pATM (S1981). Cells were untreated or treated wit 1μM olaparib for 1, 3 or 6 h, or exposed to 10 Gy of ionizing radiation. Colors indicate <5 vs ≥5 foci per marker or double-positive status.

(F) THP-1: untreated or 1, 3 and 6 h after 1μM olaparib treatment. Layout as in D. ≥ 200 cells were quantified per condition per replicate. Black dots represent the per-experiment medians (n = 3). Bars represent mean ± SD. Scalebar, 10 μm.

(B,C,F) Brown-Forsythe and Welch's one-way ANOVA with Dunnett's T3 multiple comparisons test. (D) Unpaired Welch's t-test. Indicated comparisons display numerical p-values.

Figure S2:

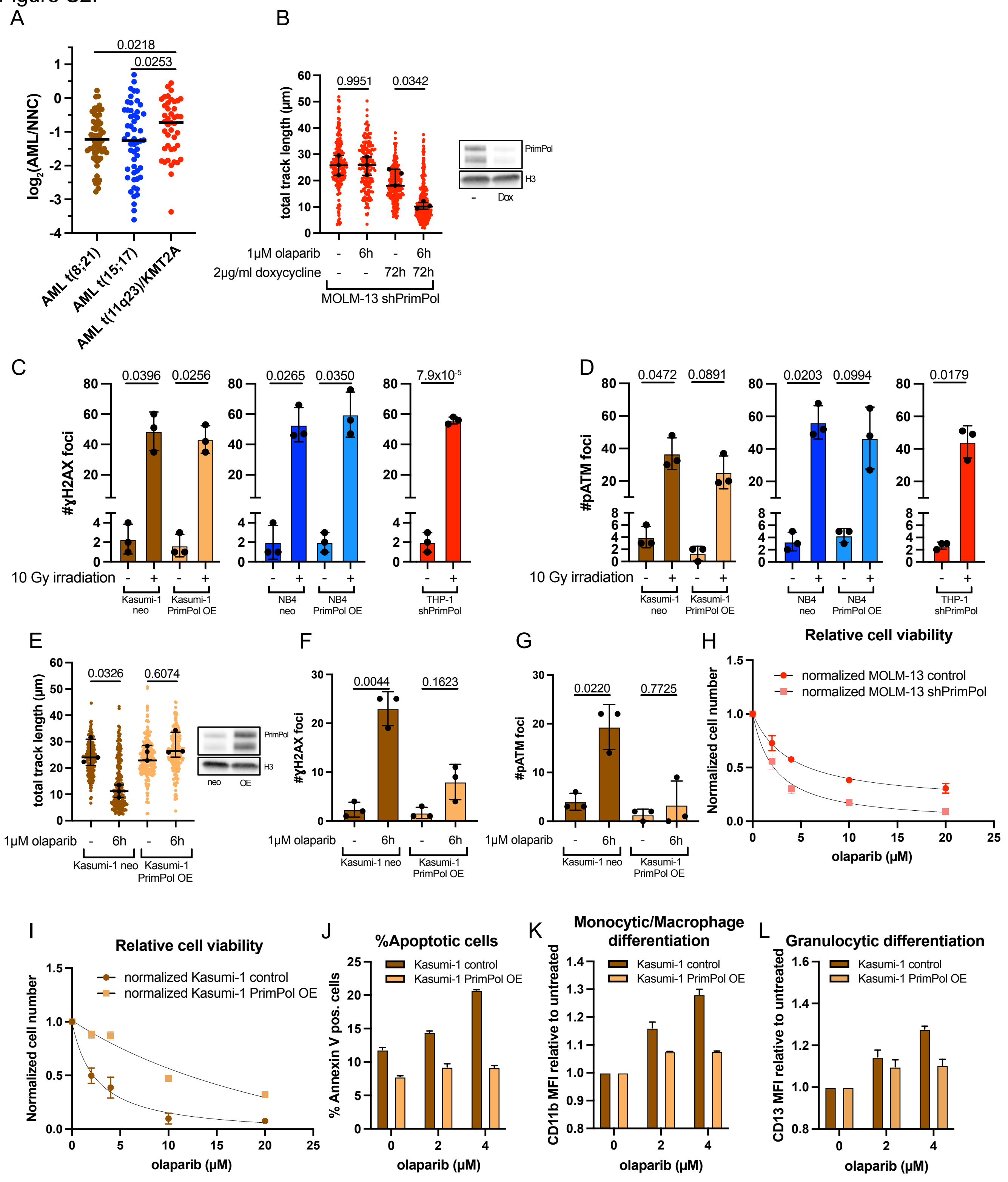
A
0.0218
0.0253
log2(AML/NNC)
AML t(8;21)
AML t(15;17)
AML t(11q23)/KMT2A
B
0.9951
0.0342
total track length (µm)
1µM olaparib
2µg/ml doxycycline
MOLM-13 shPrimPol
PrimPol
H3
Dox
C
0.0396
0.0256
0.0265
0.0350
7.9x10-5
#γH2AX foci
10 Gy irradiation
Kasumi-1 neo
Kasumi-1 PrimPol OE
NB4 neo
NB4 PrimPol OE
THP-1 shPrimPol
D
0.0472
0.0891
0.0203
0.0994
0.0179
#pATM foci
E
0.0326
0.6074
total track length (µm)
1µM olaparib
Kasumi-1 neo
Kasumi-1 PrimPol OE
PrimPol
H3
neo
OE
F
0.0044
0.1623
#γH2AX foci
G
0.0220
0.7725
#pATM foci
H
Relative cell viability
normalized MOLM-13 control
normalized MOLM-13 shPrimPol
Normalized cell number
olaparib (µM)
I
Relative cell viability
normalized Kasumi-1 control
normalized Kasumi-1 PrimPol OE
Normalized cell number
olaparib (µM)
J
%Apoptotic cells
Kasumi-1 control
Kasumi-1 PrimPol OE
% Annexin V pos. cells
olaparib (µM)
K
Monocytic/Macrophage differentiation
CD11b MFI relative to untreated
L
Granulocytic differentiation
CD13 MFI relative to untreated

**Figure S2** (related to Figure 3).

(A) PrimPol expression in AML subtypes (t(8;21), t(15;17), t(11q23)/KMT2A) was retrieved from Bloodspot (dataset "AML vs normal", GSE6891), where each AML sample's expression is compared to its nearest normal hematopoietic counterpart (NNC) as previously defined (Rapin et al., 2014). Fold-changes were computed from microarray data. We plotted $log_2$(AML/NNC).

(B) MOLM-13 with doxycycline-inducible shPrimPol: Total fiber track lengths (left) for a representative biological replicate ± 1 µM olaparib (6 h) and ± doxycycline (72 h). ≥100 fibers were scored per condition per replicate. Black line marks the median. PrimPol immunoblot showing PrimPol depletion with H3 loading control (right). Overlaid black dots represent the per-experiment median from n = 3 independent experiments; error bars = SD of those medians.

(C) ɣH2AX (Ser139) S-phase foci counts in Kasumi-1, Kasumi-1 PrimPol OE, NB4, NB4 PrimPol OE and THP-1 shPrimPol ± 10 Gy ionizing radiation. ≥200 cells per condition per replicate. Black dots represent the per-experiment medians (n = 3). Bars represent mean ± SD.

(D) pATM (S1981) S-phase foci counts in the same cell lines and irradiation as in (C). ≥200 per condition per replicate. Black dots represent the per-experiment medians (n = 3). Bars represent mean ± SD.

(E) Kasumi-1 ± PrimPol overexpression (PrimPol OE): Total fiber track lengths (left) for a representative biological replicate ± 1 µM olaparib (6 h). ≥100 fibers were scored per condition per replicate. Black line marks the median. PrimPol immunoblot confirming overexpression; H3 loading control (right). Overlaid black dots represent the per-experiment median from n = 3 independent experiments; error bars = SD of those medians.

(F) ɣH2AX (Ser139) S-phase foci counts in Kasumi-1 and Kasumi-1 PrimPol OE ± 1 µM olaparib (6h). ≥200 cells per condition per replicate. Black dots represent the per-experiment medians (n = 3). Bars represent mean ± SD.

(G) pATM (S1981) S-phase foci counts in Kasumi-1 and Kasumi-1 PrimPol OE, as in (F). ≥200 per condition per replicate. Black dots represent the per-experiment medians (n = 3). Bars represent mean ± SD.

(H) Cell-count dose-response curves for olaparib (0-20µM, 3 days) in Kasumi-1 WT and Kasumi-1 PrimPol overexpressing (PrimPol OE) cells.

(I) Annexin V-positive fraction after 3 days of olaparib (0-4µM) in Kasumi-1 WT and Kasumi-1 PrimPol OE.

(J) CD11b mean fluorescence intensity (MFI), relative to untreated, after 3 days of olaparib (0-4µM) in Kasumi-1 WT and Kasumi-1 PrimPol OE.

(K) CD13 MFI, relative to untreated, after 3 days of olaparib (0-4µM) in Kasumi-1 WT and Kasumi-1 PrimPol OE.

(L) Cell-count dose-response curves for olaparib (0-20µM, 5 days) in MOLM-13 WT and MOLM-13 doxycycline-inducible shPrimPol (MOLM-13 shPrimPol) cells.

(A-G) Brown-Forsythe and Welch's one-way ANOVA with Dunnett's T3 multiple comparisons test. (C, D) Unpaired Welch's t-test for THP-1 comparisons. Indicated comparisons display numerical p-values. (H-L) Unless indicated otherwise, bars and dose-response curve symbols indicate mean ± SD of n = 2 biological replicates.

Figure S3:

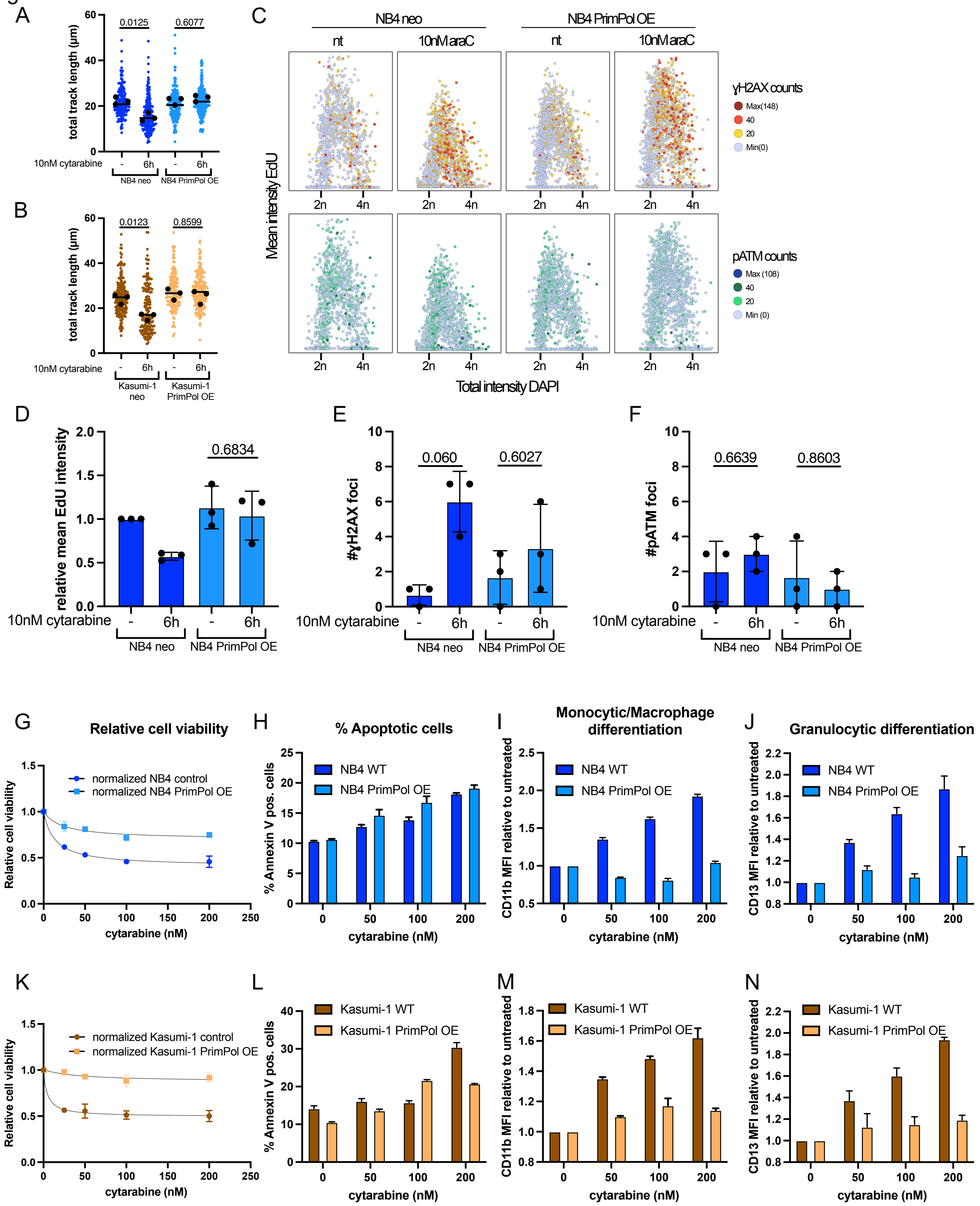
A
0.0125
0.6077
total track length (µm)
10nM cytarabine
-
6h
NB4 neo
NB4 PrimPol OE
B
0.0123
0.8599
total track length (µm)
10nM cytarabine
Kasumi-1 neo
Kasumi-1 PrimPol OE
C
NB4 neo
NB4 PrimPol OE
nt
10nM araC
γH2AX counts
Max(148)
40
20
Min(0)
pATM counts
Max (108)
Min (0)
Mean intensity EdU
Total intensity DAPI
2n
4n
D
relative mean EdU intensity
0.6834
E
#γH2AX foci
0.060
0.6027
F
#pATM foci
0.6639
0.8603
G
Relative cell viability
normalized NB4 control
normalized NB4 PrimPol OE
cytarabine (nM)
H
% Apoptotic cells
% Annexin V pos. cells
NB4 WT
NB4 PrimPol OE
I
Monocytic/Macrophage differentiation
CD11b MFI relative to untreated
J
Granulocytic differentiation
CD13 MFI relative to untreated
K
normalized Kasumi-1 control
normalized Kasumi-1 PrimPol OE
L
Kasumi-1 WT
Kasumi-1 PrimPol OE
M
N

**Figure S3** (related to Figure 3).

(A) NB4 ± PrimPol overexpression (PrimPol OE): Total fiber track lengths for a representative biological replicate ± 10 nM cytarabine (6 h). ≥100 fibers were scored per condition per replicate. Black line marks the median. Overlaid black dots represent the per-experiment median from n = 3 independent experiments; error bars = SD of those medians.

(B) Kasumi-1 ± PrimPol OE: Total fiber track lengths as in (A) ± 10 nM cytarabine (6 h). ≥100 fibers were scored per condition per replicate. Black line marks the median. Overlaid black dots represent the per-experiment median from n = 3 independent experiments; error bars = SD of those medians.

(C) NB4 ± PrimPol OE: Quantitative image-based cytometry (QIBC) cell cycle profiles based on DNA content (total DAPI intensity) and DNA synthesis (mean EdU intensity). Cells were untreated or 6 h after 10nM cytarabine (araC) treatment. Per-cell ɣH2AX (top) and pATM (bottom) foci counts are encoded by color.

(D) Relative mean EdU intensity of S-phase NB4 and NB4 PrimPol OE cells, ± 10 nM cytarabine (6h). ≥200 per condition per replicate. Black dots represent the per-experiment medians (n = 3). Bars represent mean ± SD.

(E) ɣH2AX (Ser139) S-phase foci counts in NB4 and NB4 PrimPol OE, as in (D). ≥200 cells per condition per replicate. Black dots represent the per-experiment medians (n = 3). Bars represent mean ± SD.

(F) pATM (S1981) S-phase foci counts in NB4 and NB4 PrimPol OE, as in (D). ≥200 per condition per replicate. Black dots represent the per-experiment medians (n = 3). Bars represent mean ± SD.

(G) Cell-count dose-response curves for cytarabine (0-200nM, 3 days) in NB4 WT and NB4 PrimPol OE.

(H) Annexin V-positive fraction after 3 days of cytarabine (0-200nM) in NB4 WT and NB4 PrimPol OE.

(I) CD11b mean fluorescence intensity (MFI), relative to untreated, after 3 days of cytarabine (0-200nM) in NB4 WT and NB4 PrimPol OE.

(J) CD13 MFI, relative to untreated, after 3 days of cytarabine (0-200nM) in NB4 WT and NB4 PrimPol OE.

(K) Cell-count dose-response curves for cytarabine (0-200nM, 3 days) in Kasumi-1 WT and Kasumi-1 PrimPol OE.

(L) Annexin V-positive fraction after 3 days of cytarabine (0-200nM) in Kasumi-1 WT and Kasumi-1 PrimPol OE.

(M) CD11b mean fluorescence intensity (MFI), relative to untreated, after 3 days of cytarabine (0-200nM) in Kasumi-1 WT and Kasumi-1 PrimPol OE.

(N) CD13 MFI, relative to untreated, after 3 days of cytarabine (0-200nM) in Kasumi-1 WT and Kasumi-1 PrimPol OE.

(A, B, D-F) Brown-Forsythe and Welch's one-way ANOVA with Dunnett's T3 multiple comparisons test. Indicated comparisons display numerical p-values. (G-N) Unless indicated otherwise, bars and dose-response curve symbols indicate mean ± SD of n = 2 biological replicates.
